# FusionPath: Gene fusion pathogenicity prediction using protein structural data and contextual protein embeddings

**DOI:** 10.64898/2026.01.24.701525

**Authors:** Nadine S. Kurz, Irem Berna Güven, Tim Beißbarth, Jürgen Dönitz

## Abstract

Accurate prediction of gene fusion pathogenicity is critical for understanding oncogenic mechanisms and advancing precision oncology. While existing computational methods provide valuable insights, their performance remains limited by incomplete integration of multi-scale biological features and insufficient model interpretability for clinical translation. We present FusionPath, a novel deep learning framework for gene fusion pathogenicity prediction. FusionPath uniquely integrates embeddings from multiple pretrained protein language models, including FusON-pLM and ProtBERT, alongside retained protein domains and Gene Ontology (GO) functional annotations. The model was trained and validated on a large-scale dataset of annotated pathogenic and benign fusions. The model was trained and validated on a rigorously curated dataset of 100,433 gene fusions (78,115 benign, 22,318 pathogenic) derived from FusionPDB, ChimerDB4.0, and 27 RNA-seq datasets of normal tissues. FusionPath significantly outperformed state-of-the-art methods, achieving higher AUC scores of 0.95 and 0.87 on independent test sets. By synergistically leveraging sequence, structural, and functional information with explicit modeling of wild-type sequence context, FusionPath establishes a new standard for gene fusion pathogenicity prediction by effectively leveraging complementary sequence, structural, and functional information.

## 1 Introduction

Gene fusions represent structural genomic rearrangements yielding chimeric transcripts from two previously distinct protein-coding genes. Identified in ∼17% of all cancers [1], they are a well-established class of oncogenic drivers across diverse cancers [2, 3], including *BCR-ABL1* in chronic myeloid leukemia, and *EML4-ALK* in non-small cell lung carcinoma. Critically, the clinical success of kinase-inhibiting therapies targeting fusion oncoproteins, such as imatinib for *BCR-ABL1* [4], has established them as promising targets in precision medicine [5]. Comprehensive reviews have documented the expanding landscape of targetable oncogenic fusions across diverse cancer types, highlighting their established role as therapeutic targets in precision oncology [6]. However, only a subset of these fusions are pathogenic, while others represent passenger events or occur in normal physiology [7].

Recent studies reveal that fusions tend to preserve protein function and contain specific domain architectures [8], while over half of fusion oncoproteins form biomolecular condensates with distinct functional consequences for oncogenesis [3]. These findings underscore the critical need for pathogenicity prediction frameworks that integrate protein structural and functional context to enable precision oncology applications. Furthermore, fusion oncoproteins have been associated with additional characteristics, such as RNA polymerase II partitioning [9].

Various databases have aggregated data on NGS-detected gene fusions in pan-cancer and normal tissue, including FusionGDB [10] and FusionPDB [11], and the KuNG FU (KiNase Gene FUsion) database [5]. Fusion annotation tools, including FusionAnnotator, annoFuse [12] and FusionInspector [13] focus on structural validation and basic feature annotation, but lack explicit pathogenicity scoring. Computational methods for gene fusion pathogenicity prediction include 1) ChimerDriver [14], a Multi-Layer Perceptron (MLP) integrating transcriptional and post-transcriptional information including microRNAs for oncogenic potential classification, 2) DEEPrior [15], a model combining Convolutional Neural Networks (CNN) and bidirectional Long-Short Term Memory (LSTM), 3) Pegasus [16], an approach using gradient tree boosting for binary classification of oncogenic potential, and OncoFuse [17], a Naive Bayes Network Classifier that predicts the oncogenic potential of fusion genes. Despite these advances, current methods remain limited by incomplete integration of multi-scale biological features and lack of interpretability, highlighting the need for more sophisticated approaches that can capture the complex molecular determinants of fusion pathogenicity [18].

We present FusionPath, an interpretable framework for gene fusion pathogenicity prediction. FusionPath integrates three complementary biological scales, including fusion-specific protein embeddings via language models, domain architecture alterations, and Gene Ontology (GO) [19] term enrichment. By unifying sequence-level, structural, and systems-level features, FusionPath provides biologically grounded pathogenicity scores with mechanistic insights.

## 2 Methods

### 2.1 Datasets

We compiled gene fusion data from two primary sources: FusionPDB, a structural database of experimentally validated oncogenic fusions, and ChimerDB4.0 [20], which integrates literature-mined, cancer-type-specific fusions. To retrieve additional gene fusions from normal tissues, we utilized 27 RNA-seq datasets of the European Nucleotide Archive (ENA), including fusions from 95 healthy donors of 27 tissues were downloaded from PRJEB4337 [21], and fusions from cancerous and normal breast cancer tissue were downloaded from PRJNA975550 [22], among others. The full list of datasets is available at Supplementary Table S1.

### 2.2 Preprocessing and annotation

All genomic coordinates were standardized to GRCh38 using the SeqCAT LiftOver API [23]. Fusion calling for normal tissues was performed with STARFusion [24, 25], yielding 5.403 fusion candidates. Wild-type protein sequences of the fusions’ head and tail genes were queried using SeqCAT’s protein sequence API endpoint, retrieving the genes’ MANE Select transcripts [26]. Protein domain and GO terms were retrieved using InterProScan 5 [27]. Known gene roles of tumor suppressor and oncogenes were retrieved from Onkopus [28] gene role Application Programming Interface (API) endpoint, where we selected the classification by Vogelstein et al. (2013) [29]. Gene-disease associations were retrieved from Clingen [30].

### 2.3 The FusionPath model

For each fusion protein sequence, FusionPath extracts the wild-type protein sequences of the affected genes, as well as the retained protein domains and GO terms identified in the fusion sequence. The fusion amino acid sequence was encoded using FusON-pLM [31]. Wild-type head/tail sequences were independently embedded via Prot-BERT [32]. Concatenated sequence embeddings underwent batch normalization and dropout. Concurrently, retained protein domains and Gene Ontology terms were tokenized into a vocabulary, projected to 128-dim embeddings, and processed via additive attention [33]. To dynamically weight the contribution of each domain and GO term based on its relevance to pathogenicity, we employ an additive attention mechanism. Given an embedding matrix **E** ∈ ℝ^*x*×*d*^ containing the d-dimensional embeddings of n retained protein domains and GO terms, the attention weights are computed as:

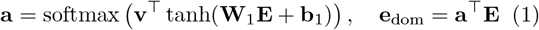

where **W**_1_ ∈ ℝ^*h*×*d*^ is a learnable weight matrix, *b*_1_ ∈ ℝ^*h*^ is a bias term, **v** ∈ ℝ^*h*^ is a context vector, and *h* is the hidden dimension. The processed sequence embeddings and functional annotation embeddings are then concatenated into a unified feature vector **z** ∈ ℝ^*D*^, where *D* is the total dimensionality of the integrated representation. Integrated features fed a 3-layer Multi-layer perceptron (MLP), with sigmoid output for pathogenicity probability.

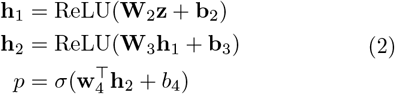

where 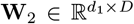 and 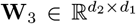are weight matrices for the hidden layers with dimensions *d*_1_ = 512 and *d*_2_ = 256, **b**_2_ and **b**_3_ are corresponding bias vectors, 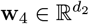 is the output layer weight vector, *b*_4_ is the output bias term, and *σ*(·) denotes the sigmoid function. The model was trained as a binary classifier, optimizing binary cross-entropy loss with logits (BCEWithLogitsLoss) using the AdamW optimizer. Given *N* training samples with true labels *y*_*i*_ ∈ {0, 1} and raw model outputs (logits) *z*_*i*_, the loss function is defined as:

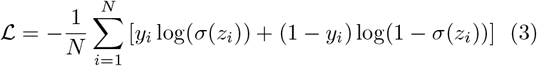

where *z*_*i*_ are the raw logits. Training proceeded with standard forward-backward passes, where the model outputs were passed through a sigmoid function.

### 2.4 Evaluation metrics

We employed receiver operating characteristic (ROC) curve analysis and the area under the ROC curve (AUC) for evaluating the model’s performance compared to prior methods. The impact of protein domains and GO terms was assessed by computing attention weights. SHAP (SHapley Additive exPlanations) [34] values were computed using the KernelExplainer with 500 background samples drawn from the test dataset, with 2000 Monte Carlo iterations for precision. Attention weights were computed using an additive attention mechanism [33], where feature importance was quantified as the mean attention weight across high-probability pathogenic samples. Only features appearing in ≥5 samples were included in the final ranking, with results sorted by mean attention weight to identify the most influential protein domains and GO terms.

## 3 Results

### 3.1 The FusionPath architecture and dataset

FusionPath implements a multimodal model that combines protein structural information with the contextual embeddings of pretrained protein language models. The model processes amino acid sequences of fusion proteins as input and extracts the wild-type sequences of the head and tail genes, as well as the retained protein domains and GO terms (Figure 1). The model incorporates two specialized components: 1) FusON-pLM, a protein language model fine-tuned exclusively on oncoprotein sequences to process fusion protein contexts, and 2) ProtBERT, used to extract implicit functional knowledge from wild-type sequences. In addition, the model processes the retained protein domains as well as identified GO terms as input.

**Figure 1:**
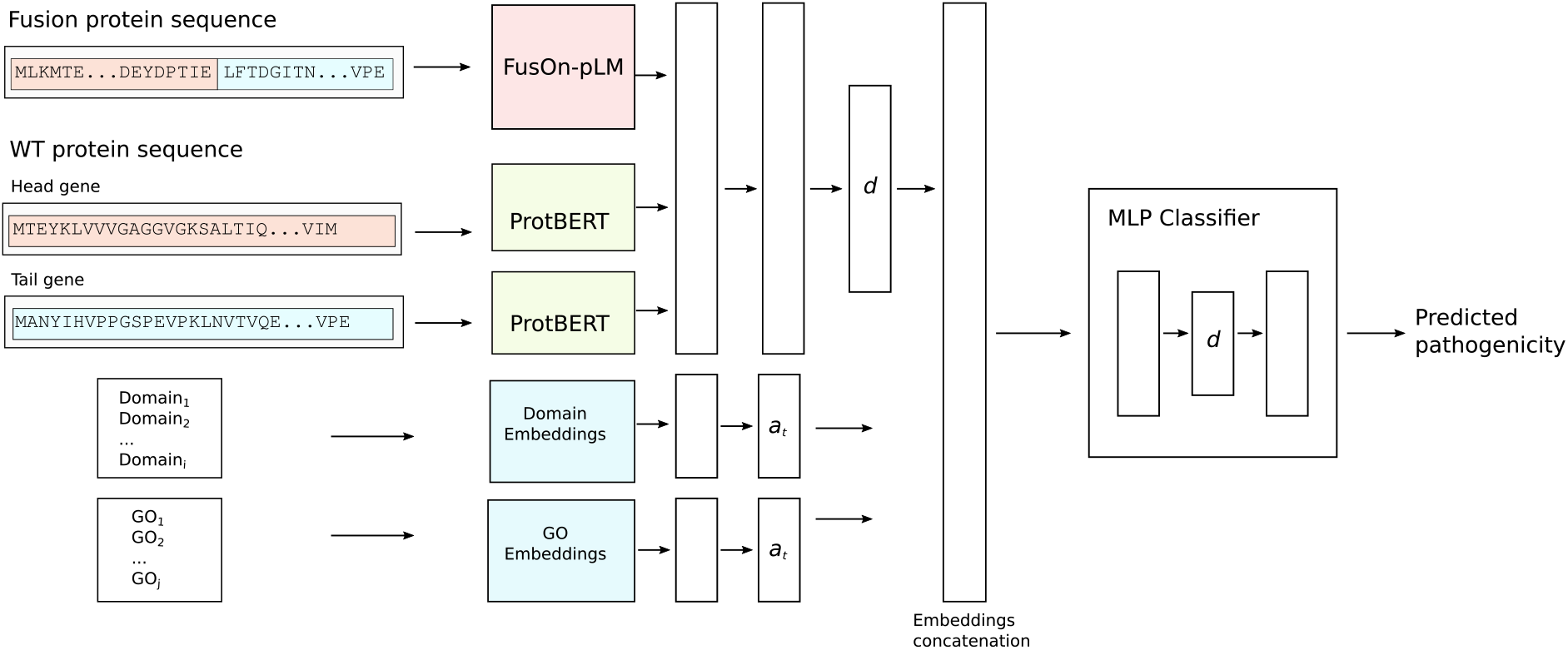
The FusionPath architecture. The fusion amino acid sequence is processed by FusON-pLM, generating contextual embeddings. The wild-type head and tail gene sequences are independently encoded using ProtBERT, yielding separate embedding tensors for each sequence. Embeddings from all sequence streams are concatenated along the feature dimension, followed by batch normalization and dropout regularization. Protein domains and and GO terms associated with the fusion are mapped to discrete tokens using a curated biological vocabulary. Tokenized inputs are then projected into a dense embeddings and processed through a dedicated attention layer, enabling context-aware weighting of functionally relevant domains and GO annotations. The processed sequence embeddings and functional annotation embeddings are then concatenated into a unified feature vector. This integrated representation is passed through a multilayer perceptron (MLP) classifier comprising ReLU activation and dropout. The final layer uses a sigmoid activation to output a pathogenicity probability score.

We curated a dataset of 100.433 gene fusions from cancerous and normal tissue databases, supplemented by 27 manually validated normal tissue RNA-seq datasets. For classifying gene fusions, we annotated the head and tail genes of the fusions with their known role in carcinogenesis (tumor suppressors and oncogenes), known genedisease associations. Fusions were labeled as pathogenic if they were associated with cancer, involved a known oncogene or tumor suppressor, or a gene-disease association. Fusions were labeled as benign if they did not include a known tumor suppressor or oncogene, had no known genedisease association, and whose fusion protein as well as the head or tail gene had no known association as cancer biomarker. Additionally, all fusions that were called from normal tissue were labeled as benign. The final dataset comprised 78.115 benign (77.8%) and 22.318 pathogenic (22.2%) fusions.

### 3.2 FusionPath improves pan-cancer gene fusion pathogenicity prediction

We benchmarked FusionPath against the three state-ofthe-art methods ChimerDriver, DEEPrior, and Pegasus. Using our curated dataset of 19.934 validated pathogenic and benign gene fusions, we assessed the model performance. FusionPath achieved significantly higher prediction accuracy compared to the other methods, with an AUC of 0.95% on the test data set (Figure 2A). To assess generalizability, we further evaluated FusionPath on the DEEPrior training dataset as an external validation cohort, achieving an AUC of 0.87 (Figure 2B), confirming robustness across diverse data sources.

**Figure 2.**
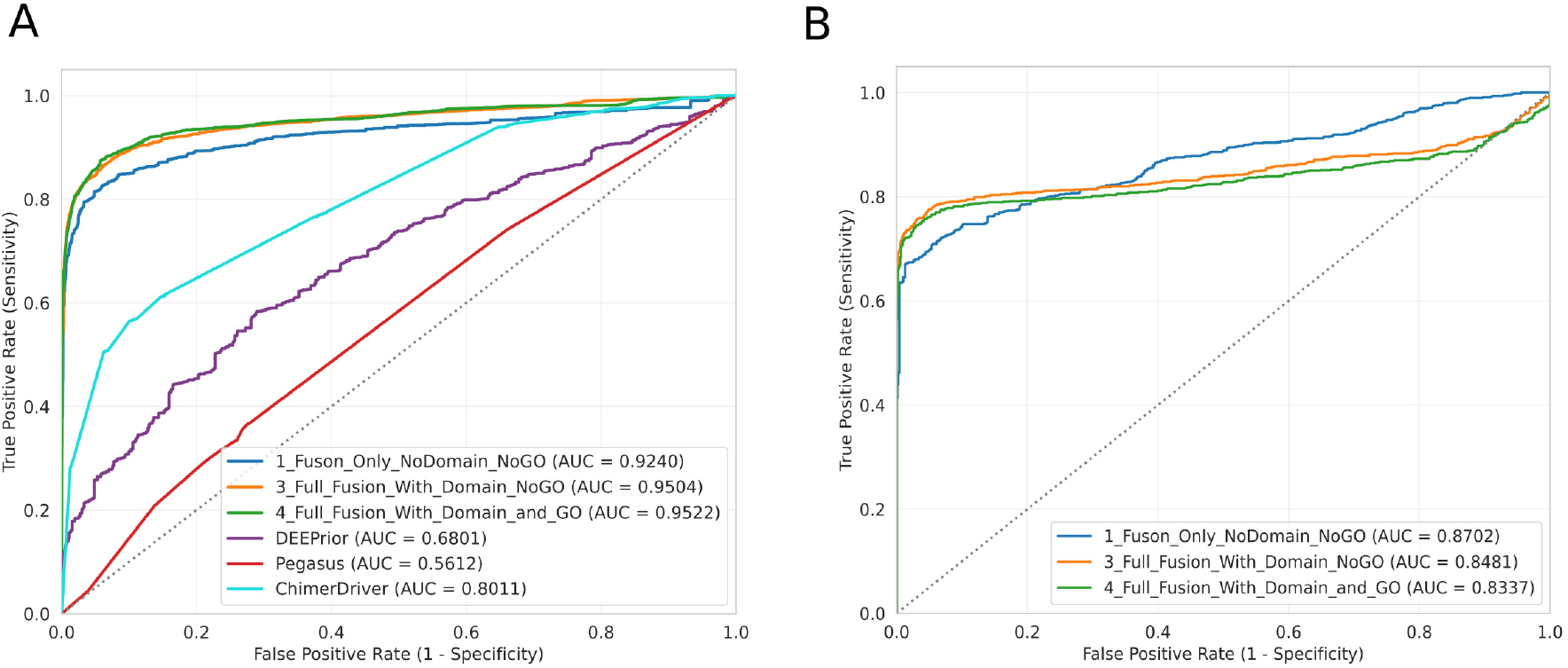
Performance comparison and classification results of FusionPaths.(**A**) ROC/AUC curves of different models of FusionPath on the test set containing 19.934 fusion protein, including models with and without the inclusion of the Prot-BERT models wild-type sequence embeddings, as well as with and without the inclusion of protein structure. The full model (integrating wild-type sequence embeddings, protein structure, domains, and GO terms) achieved the highest AUC (0.95). All models significantly outperformed ChimerDriver (AUC=0.8011), DEEPrior (AUC=0.68), and Pegasus (AUC=0.56). (**B**) ROC/AUC curve of FusionPath tested on the DEEPrior training dataset, achieving an AUC of 0.87.

### 3.3 FusionPath provides insights in the role of protein domains and functions in fusion proteins

Beyond predictive accuracy, FusionPath provides biologically interpretable insights through SHAP analysis as well as its attention mechanism, which quantify feature contributions, including the protein domains and GO terms, to pathogenicity predictions. This enables identification of molecular determinants driving oncogenicity, facilitating hypothesis generation for experimental validation. Analysis of the test set revealed cancer-typespecific pathogenicity patterns, with the highest average pathogenicity in Acute Myeloid Leukemia (LAML) and Mesothelioma (MESO), and the lowest in Adrenocortical Carcinoma (ACC), Kidney Renal Clear Cell Carcinoma (KIRC), and Uveal Melanoma (UVM) (Figure 3A). Critically, SHAP analysis identified protein kinase domains (e.g., serine-threonine/tyrosine kinases) as the strongest positive contributors to pathogenicity (Figure 3B), consistent with known oncogenic mechanisms like constitutive kinase activation. Conversely, domains such as fibroblast growth factor receptors (FGFRs) and immunoglobulin-like domains exhibited the most negative SHAP values, suggesting protective or neutral roles in fusion contexts. This domain-level resolution reveals how specific functional modules dictate fusion-driven oncogenesis.

**Figure 3.**
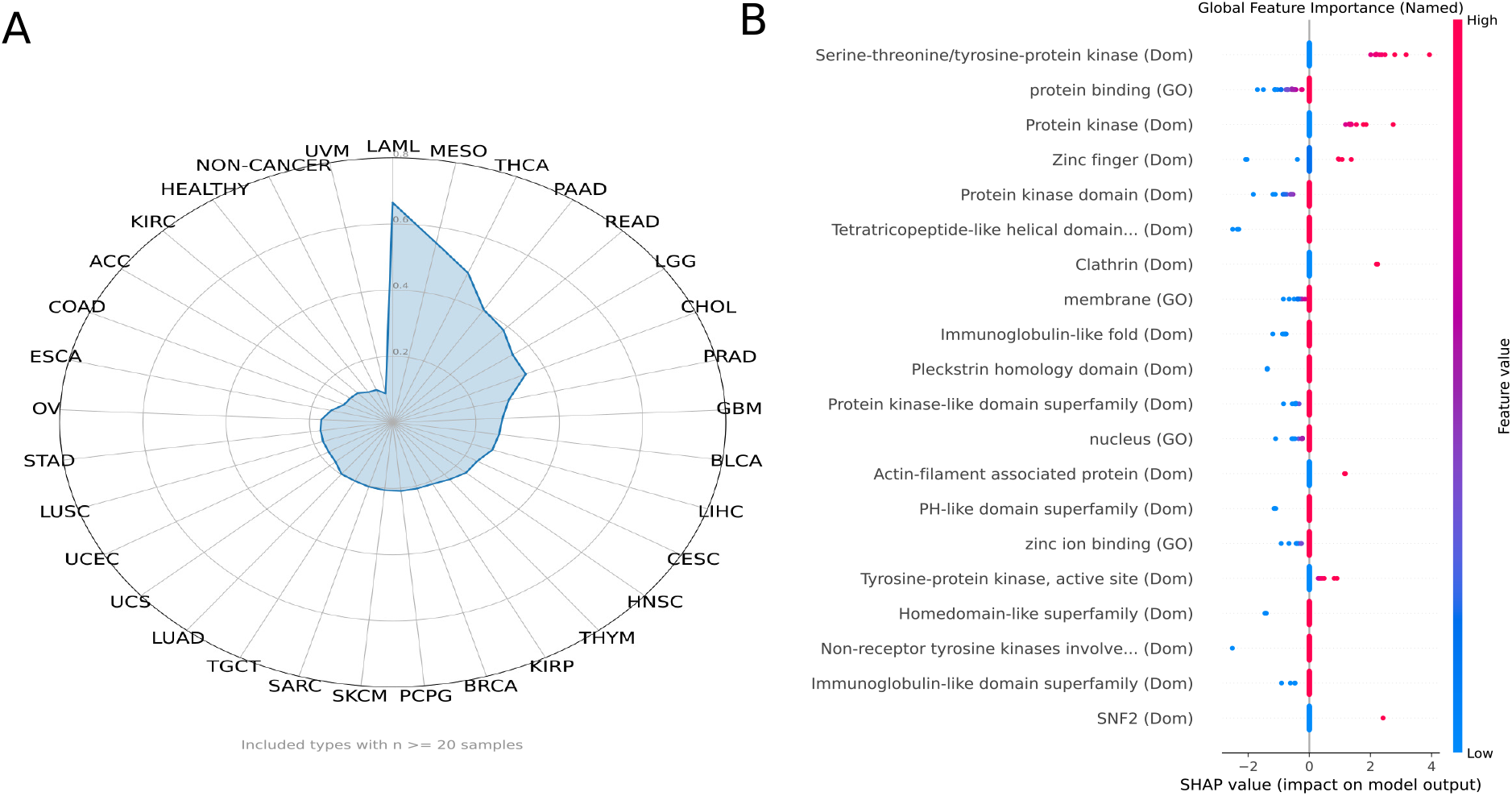
Analysis of the effect of protein domains and GO terms on gene fusion pathogenicity.(**A**) Average predicted gene fusion pathogenicity per cancer type. Acute Myeloid Leukemia (LAML), Mesothelioma (MESO) and Thyroid carcinoma (THCA) exhibited the highest average fusion protein pathogenicity, while the most benign fusions could be identified in Adrenocortical carcinoma (ACC), kidney renal clear cell carcinoma (KIRC), and uveal melanoma (UVM). (**B**) SHAP values for FusionPath detecting how individual protein domains and GO terms shift the model’s prediction. Positive SHAP values indicate that a domain/GO term increases predicted pathogenicity, while negative values indicate a decrease.

### 4 Discussion

Accurate identification of pathogenic gene fusions remains a critical need in precision oncology, where misclassification may lead to missed driver mutations, ineffective therapies or overlooked therapeutic opportunities. We present FusionPath, a multimodal framework integrating protein language modeling with functional genomic context to predict fusion pathogenicity. Our results demonstrate that FusionPath significantly outperforms state-of-the-art methods in both pan-cancer and tissue-specific benchmarks. FusionPath’s integration of protein structural data with contextual embeddings and functional annotations enables more comprehensive assessment across diverse fusion types, addressing a key limitation identified in the benchmark where tools struggled with fusions lacking obvious cancer-related gene associations [18]. This capability is particularly valuable for discovering novel fusion drivers beyond the well-characterized kinase fusions. FusionPath’s performance stems from its integration of complementary biological signals through three synergistic components: A FusON-pLM model embedding the chimeric protein sequence to capture novel structural and functional features induced by the fusion junction, dual ProtBERT encoders processing the wild-type parental sequences to preserve baseline functional context disrupted by the fusion, and layers integrating protein domain architectures and GO terms to explicitly model functional consequences, e.g. loss of regulatory domains, or gain of dimerization motifs.

A major challenge in predicting the pathogenicity of gene fusions is determining the ground truth dataset that serves as the basis for training and validating the model. While two human-curated datasets, ClinVar [35] and HGMD [36], are available for missense variants and are often employed for model validation for missense variant pathogenicity prediction, there is no comparable equivalent for gene fusions or structural variants. This challenge was starkly demonstrated by Wei et al., who systematically re-evaluated 2,727 reported prostate cancer fusions using TCGA RNA-seq data and found that over 90% could not be validated in independent cohorts, with only 4% showing tumor-specific occurrence [37]. Existing fusion databases, e.g. COSMIC [38], catalog occurrences, but lack systematic pathogenicity labels. To overcome this, we constructed a rigorously curated dataset of cancer-derived and normal-tissue gene fusions, where fusions were labeled pathogenic if they involved a known tumor suppressor gene or oncogene, included at least one gene disease association. This approach corresponds to biological reality, as most gene fusions represent passenger rather than driver mutations. However, a critical consideration is the class imbalance in the dataset (77.8% benign fusions), which inherently biases models toward benign predictions. While our biologically informed curation criteria (cancer recurrence, tumor suppressor gene/oncogene involvement, disease association) provide a pragmatic and interpretable proxy for pathogenicity, it has to be pointed out that this approach does not imply with certainty that all fusions are correctly classified.

Notably, FusionPath revealed cancer-type-specific pathogenicity patterns, with LAML, MESO, and THCA exhibiting the highest scores, and ACC, KIRC, and UVM showing the lowest. While requiring experimental validation, these patterns align with known oncogenic mechanisms. For instance, high scores in THCA reflect the established role of RET and NTRK fusions in subsets of papillary thyroid carcinoma, which are actionable targets for kinase inhibitors [cite specific reviews, e.g., Cancer Discov 2018]. As these are tumor types with fusions established as therapeutic targets, e.g. *BCR::ABL1, RUNX1::RUNX1T1* or *ETV6::NTRK3* in LAML [39, 40], FusionPath has direct clinical relevance and can contribute to improved targeted therapies for fusion oncoproteins. Conversely, the low average pathogenicity in UVM is consistent with its primary dependence on GNAQ/GNA11 mutations rather than fusion drivers [41].

SHAP interpretability analysis revealed kinase-related domains, including receptor tyrosine kinases, fibroblast growth factor receptors, and serine-threonine/tyrosine kinases, as the dominant predictors of pathogenicity, directly aligning with established oncogenic mechanisms where fusion-driven kinase activation drives malignancy. Critically, the model prioritizes therapeutically targetable kinase fusions (e.g. FGFR, ALK) with high confidence, enabling direct translation to biomarker-guided therapy selection. Despite known database overrepresentation of kinase fusions, rigorous validation on kinase-depleted cohorts confirmed robust generalizability to non-kinase pathogenic events. This kinase-centric interpretability establishes our tool as a biologically grounded clinical decision aid for precision oncology.

In summary, FusionPath establishes a new paradigm for fusion pathogenicity prediction by unifying deep protein language modeling with explicit functional genomics. By providing a transparent, biologically grounded dataset and a model that reveals fusion protein pathogenicity, we offer a tool that can accelerate both discovery of novel fusion drivers and clinical decision-making in precision oncology.

## Code and data availability

The source code of FusionPath is available at https://gitlab.gwdg.de/MedBioinf/mtb/fusionpath.

## Author contributions

NSK conceptualized the method. NSK and IBG implemented the model. NSK wrote the manuscript. All authors proofread the manuscript.

## Acknowledgments

The authors thank the Internal Max Planck Research School for Genome Science (IMPRS-GS).

